# Characterization of newly isolated bacteriophages targeting carbapenem-resistant *Klebsiella pneumoniae*

**DOI:** 10.1101/2024.09.29.615722

**Authors:** Bokyung Kim, Shukho Kim, Yoon-Jung Choi, Minsang Shin, Jungmin Kim

**Author notes:** These authors contributed equally to this work (Bokyung Kim, Shukho Kim). Corresponding Author’s information To whom correspondence should be addressed. **Address** : Gukchaebosang-ro 680, Jung-gu, Daegu 41944, Republic of Korea **(E-mail)**.

## Abstract

*Klebsiella pneumoniae*, a Gram-negative opportunistic pathogen, is increasingly resistant to carbapenems in clinical settings. This growing problem necessitates the development of alternative antibiotics, with phage therapy being one promising option. In this study, we investigated novel phages targeting carbapenem-resistant *Klebsiella pneumoniae* (CRKP) and evaluated their lytic capacity against clinical isolates of CRKP. First, 23 CRKP clinical isolates were characterized using Multi-locus Sequence Typing (MLST), carbapenemase test, string test, and capsule typing. MLST classified the 23 *K. pneumoniae* isolates into 10 sequence types (STs), with the capsule types divided into nine known and one unknown type. From sewage samples collected from a tertiary hospital, 38 phages were isolated. Phenotypic and genotypic characterization of these phages was performed using Random Amplification of Polymorphic DNA-PCR (RAPD-PCR), transmission electron microscopy (TEM), and whole genome sequencing (WGS) analysis. Host spectrum analysis revealed that each phage selectively lysed strains sharing the same STs as their hosts, indicating ST-specific activity. These phages were subtyped based on their host spectrum and RAPD-PCR, identifying nine and five groups, respectively. Fourteen phages were selected for further analysis using TEM and WGS, revealing 13 *Myoviruses* and one *Podovirus*. Genomic analysis grouped the phages into three clusters: one closely related to *Alcyoneusvirus*, one to *Autographiviridae*, and others to *Straboviridae*. Our results showed that the host spectrum of *K. pneumoniae*-specific phages corresponds to the STs of the host strain. These 14 novel phages also hold promise as valuable resources for phage therapy against CRKP.

## Introduction

*Klebsiella pneumoniae* is a Gram-negative, encapsulated, and opportunistic bacterium capable of causing severe illnesses, including pneumonia, liver abscesses, urinary tract infections, and bloodstream infections. Some *K. pneumoniae* strains are highly virulent due to their production of mucoid capsules, which protect them from the host immune system and antibiotics. Hypervirulent *K. pneumoniae* (hvKP) has emerged as a significant cause of serious, life-threatening infections in healthy individuals within the community owing to its production of capsular polysaccharides. This poses a significant challenge for phage treatment (Tang et al., 2023). In clinical settings, the overuse of antibiotics has led to the emergence of multi-drug resistance (MDR) *K. pneumoniae*, including carbapenem-resistant *Klebsiella pneumoniae* (CRKP), which is a life-threatening pathogen. As carbapenems are considered the last-resort antibiotic, the problem of carbapenem resistance is becoming severe. Therefore, the development of alternative antibiotics is urgently required, and bacteriophage (phage) therapy is a strong candidate for addressing this issue. (Zurabov and Zhilenkov, 2021; Townsend et al., 2021; Baqer et al., 2022; Peng et al., 2023; Choi et al., 2024).

Phages are viruses that target and infect bacteria by attaching to specific receptors on the bacterial cell surface, establishing a potent and highly evolved host-parasite interaction. Unlike broad-spectrum antibiotics, which can impact a wide range of bacteria, phages are highly selective and only affect the bacterial strains they are designed to target. This high degree of specificity enhances their safety profile and minimizes the risk of collateral damage to beneficial bacteria in the body or the environment. Phages are prevalent and incredibly diverse, making them one of the most abundant organisms on Earth. They exist in diverse environments such as water, soil, and even within the bodies of animals. Owing to these characteristics, phage therapy, which uses phages as antibacterial agents, has fewer side effects than antibiotics and shows promise in combating MDR bacterial infections (Zurabov and Zhilenkov, 2021; Townsend et al., 2021; Baqer et al., 2022; Peng et al., 2023). Although phage therapy has been studied for a century, interest in it has recently resurged due to antibiotic resistance, exploring it as an alternative approach to treating bacterial infections (Gordillo and Barr, 2019). The rapid proliferation of antibiotic resistance due to antibiotic overuse and the notably slow pace of new antibiotic development (Nilsson, 2019) underscore the promising potential of phage therapy as an alternative. Despite its potential of phage therapy, numerous hurdles persist, including regulatory, ethical, and pharmacokinetic challenges in clinical settings. Transitioning phage therapy from theoretical potential to practical application involves establishing a well-characterized phage library. This library would consist of a diverse collection of phages, each extensively studied and categorized to determine their specificity, potency, and safety profiles. This comprehensive understanding of individual phages would facilitate the selection of the most appropriate candidates for specific bacterial infections.

We aimed to establish phage resources for therapy against *K. pneumoniae* infections by characterizing prevalent *K. pneumoniae* isolates in Korea and newly isolated phages that can lyse those bacteria. This study characterized *K. pneumoniae* clinical isolates obtained from various patients through antibiotic susceptibility tests, carbapenem resistance determination, and multilocus sequence typing (MLST). Also, we characterized 14 phages selected from 38 phage isolates by conducting host spectrum determination and genotypic and phenotypic analyses. The promising phage candidates’ corresponding subtypes of CRKP were described and discussed.

## Materials and Methods

### Bacterial Isolates

A total of 23 clinical isolates of MDR *K. pneumoniae* and CRKP were obtained from Kyungpook National University Hospital in Daegu, Korea. These isolates, collected from various clinical sources such as pus, urine, sputum, and wounds between 2011 and 2022, were analyzed for their MDR profiles using the VITEK 2 system (bioMérieux, France), which provides automated antimicrobial susceptibility testing. Relevant clinical information is detailed in Table 1. The bacteria were cultured on Blood Agar Plate (BAP; Synergy Innovation, Seongnam, Korea) at 37[for 18 h and/or in Brain Heart Infusion (BHI; Becton Dickinson and Co., Sparks, MD, USA) broth with shaking at 150 rpm at 37[for 18 h depending on the experimental requirements.

**Table 1.** Sequence types, carbapenemase types, and capsule types of carbapenem-resistant Klebsiella pneumoniae clinical isolates used in this study.

| Isolate | Isolation year | Source | ST type | CARBA-NP test | String test | wzi allele | K-type or KL-type |
| --- | --- | --- | --- | --- | --- | --- | --- |
| KBN10P02346 | 2011 | Pus | ST11 | + | - | 50 | K15,K17,K50,K51,K52 |
| KBN10P02349 | 2011 | Bile | ST336 | - | - | 141 | KL25 |
| KBN10P05309 | 2017 | Urine | ST11 | + | + | 123 | KL136 |
| KBN10P06378 | 2019 | Wound | ST307 | + | - | 173 | KL102, KL149, KL155 |
| KBN10P06492 | 2019 | Rectal swab | ST789 | + | - | 18 | K18 |
| KBN10P06644 | 2019 | Rectal swab | ST307 | + | + | 173 | KL102, KL149, KL155 |
| KBN10P06655 | 2019 | Rectal swab | ST133 | - | - | 180 | KL116 |
| KBN10P06710 | 2020 | Sputum | ST15 | + | - | 19 | K19 |
| KBN10P06778 | 2020 | Rectal swab | ST307 | + | - | 173 | KL102, KL149, KL155 |
| KBN10P06805 | 2020 | Venous blood | ST307 | + | - | 173 | KL102, KL149, KL155 |
| KBN10P07139 | 2020 | Urine | ST307 | + | - | 173 | KL102, KL149, KL155 |
| KBN10P07146 | 2020 | Bile | ST307 | + | + | 173 | KL102, KL149, KL155 |
| KBN10P07153 | 2020 | Central blood | ST307 | + | - | 173 | KL102, KL149, KL155 |
| KBN10P07294 | 2021 | Sputum | ST48 | - | - | 62 | K62 |
| KBN10P07305 | 2021 | Sputum | ST7145 | + | - | 173 | KL102, KL149, KL155 |
| KBN10P07385 | 2021 | Urine | ST307 | + | - | 173 | KL102, KL149, KL155 |
| KBN10P07391 | 2021 | Urine | ST307 | + | - | 173 | KL102, KL149, KL155 |
| KBN10P07398 | 2021 | Rectal swab | ST307 | + | - | 173 | KL102, KL149, KL155 |
| LIS20220441 | 2022 | Venous blood | ST307 | + | - | 173 | KL102, KL149, KL155 |
| LIS20220486 | 2022 | Venous blood | ST392 | + | - | 178 | Unknown |
| LIS20222037 | 2022 | Venous blood | ST11 | + | - | 123 | KL136 |
| LIS20222038 | 2022 | Central blood | ST11 | + | - | 123 | KL136 |
| LIS20222290 | 2022 | Venous blood | ST307 | + | - | 173 | KL102, KL149, KL155 |
| ATCC13883 |  |  | ST3 | - | + | 3 | K3 |

### Carbapenemase Test of CRKP

The Rapidec^®^ Carba NP test (bioMérieux, France) was conducted on 23 clinical isolates to detect carbapenemase production. The procedure followed the manufacturer’s instructions. Briefly, bacteria were cultured on BAP and colonies were taken using a sterile wood stick. Results were initially observed after a 30-min incubation and re-evaluated after 2 h.

### Carbapenemase Gene Determination of CRKP

Carbapenemase genes were screened using polymerase chain reaction (PCR) with specific primers targeting *bla*_KPC-2_, *bla*_IMP_, *bla*_VIM_, *bla*_NDM_, and *bla*_OXA-48_ (Bonnin et al., 2021). For this study, only *bla*_KPC-2_ (forward: 5’-CGTCTAGTTCTGCTGTCTTG-3’, reverse: 5’-CTTGTCATCCTTGTTAGGCG-3’) and *bla*_OXA-48_ (forward: 5’-GCGTGGTTAAGGATGAACAC-3’, reverse: 5’-CATCAAGTTCAACCCAACCG-3’) were selected for experimental analysis. PCR reactions were performed using AccuPower PCR PreMix (BIONEER, Daejeon, Korea) under the following conditions: initial denaturation at 94[for 10 min; 36 cycles of 94[for 30 s, 52[for 40 s, and 72[for 50 s; followed by a final extension at 72[for 5 min (Poirel et al., 2011).

### Capsule Typing of CRKP

The capsule types of CRKP isolates were determined through *wzi* gene sequencing. The PCR amplification and sequencing of the *wzi* gene were performed following the method described by Brisse *et al*. (2013) (Brisse et al., 2013). Amplification utilized the primers *wzi*_F (5’-GTGCCGCGAGCGCTTTCTATCTTGGTATTCC-3’) and *wzi*_R (5’-GAGAGCCACTGGTTCCAGAACTTCACCGC-3’) with AccuPower PCR PreMix. The PCR conditions were as follows: an initial denaturation at 94[for 2 min; 30 cycles of 94[for 30 s, 55[for 40 s, and 72[for 30 s, followed by a final extension at 72[for 5 min. The capsule types of 23 *K. pneumoniae* isolates were determined using the Pasteur Klebsiella database (PasteurMLST. Retrieved from https://bigsdb.pasteur.fr/cgi-bin/bigsdb/bigsdb.pl?db=pubmlst_klebsiella_isolates).

### String Test of CRKP

A string test detected hyper muco-viscosity (HMV) in the CRKP isolates. The CRKP isolates were streaked on a BAP and incubated for 18 h at 37[. The presence of HMV was confirmed if a viscous string >5 mm in length formed when a colony on the agar plate was stretched with a loop (Hagiya et al., 2014).

### MLST of CRKP

MLST was conducted to identify the clonal lineages of the bacterial isolates. Seven housekeeping genes (*gapA*, *infB*, *mdh*, *pgi*, *phoE*, *rpoB*, and *tonB*) were amplified and sequenced. The amplification conditions were as follows: one cycle at 94[for 2 min; 35 cycles of 94[for 20 s, 50[for 30 s, and 72[for 30 s; followed by a final extension at 72[for 5 min. The resulting amplicons were sequenced by Macrogen, Inc. (Seoul, Korea). MLST assignments for each isolate were generated from the sequence data, and unknown sequence types (STs) were submitted to the *K. pneumoniae* MLST database at the Pasteur Institute (PasteurMLST). The MLST scheme is described in detail in the PasteurMLST.

### Phage Isolation

Fifteen sewage samples were collected from a tertiary hospital between April 2022 and September 2022. The samples were centrifuged for 20 min at 7,000 rpm for sample preparation. The supernatant was filtered using a 0.45 μm syringe filter, chloroform was added to a final concentration of 10% (v/v), and the mixture was stored at 4[. For phage enrichment, 1 mL of the pretreated sewage sample was added to 10 mL of bacterial solution (optical density [OD] = 0.5 at 600 nm) containing KCTC 2208, KBN10P05309, ATCC13883, and KBN10P07398 as bacterial hosts. The samples were incubated in a shaking incubator at 30[for 24 h, followed by storage at 4[for 48 h. The solution was then centrifuged at 7,000 rpm for 10 min. The supernatant was filtered using a 0.22 μm syringe filter, chloroform was added to a final concentration of 10% (v/v), and the mixture was stored at 4[. The presence of lytic phages was confirmed by performing a spot assay on the bacterial lawn of the original enriching hosts (i.e., KCTC 2208, KBN10P05309, ATCC13883, and KBN10P07398). Each phage was isolated by obtaining a single plaque through a plaque assay. Pure phage propagation was conducted using 5 mL of bacterial solution (OD = 0.5 at 600 nm), 150 μL of phage solution, MgSO_4_ (final concentration 2 mM), and CaCl_2_ (final concentration 1 mM). The mixture was incubated at 37[for 18 h.

### Phage Storage

Phage concentration and buffer exchange from BHI to SM buffer (100 mM NaCl, 8.0 mM MgSO_4_, 50 mM Tris pH 7.5, and 0.001% gelatin) were performed using an Amicon Ultra-15 centrifugal filter (50 K) device (Merck Millipore Ltd.). The centrifugal filtration process increased concentrated the phage titer by 10 to 100 times in a single use. After filtration, phage stocks were prepared by adding sterile glycerol to a final concentration of 20% and stored in a deep freezer.

### Phage Plaque Assay and Spot Assay

A spot assay was performed to verify the formation of phage lytic spots. One milliliter of bacterial solution (OD 600 nm = 0.5) was mixed with 20 mL of 0.75% BHI soft agar and distributed onto a square plate. Ten microliters of each serially diluted phage solution (from 10^-1^ to 10^-10^) were then dropped onto the soft agar plate, which was incubated overnight at 37[. A plaque assay was performed to measure the phage titer and assess lytic activity and plaque morphology. Five hundred microliters of bacterial solution (OD 600 nm = 0.5), 100 μL of each serially diluted phage solution (from 10^-1^ to 10^-10^), and 10 mL of 0.75% BHI soft agar were mixed and poured into a 90-mm Petri dish. Phages were diluted with SM buffer. After incubating overnight at 37[, the plaques were counted. Petri dishes with countable plaques (30–300 plaques) were selected to evaluate the plaque forming unit (PFU).

### Host Spectrum Determination of Phages

Using the double-layer method, the lytic spectra of 38 phages were determined by spot assay against 23 clinical isolates of CRKP and one standard strain (*K. pneumoniae* ATCC 13883). A phage titer exceeding 10^10^ PFU/mL was used for the test, and 10 μL of this phage solution was spotted on each bacterial lawn.

After 18 h of incubation at 37[, the lysis zones were categorized as follows: ([) CL: clear transparent lysis zone without any resistant colonies; ([) SCL: semi-clear lysis zone with some resistant colonies; ([) OL: non-transparent opaque lysis zone. Based on the spot assays results, a plaque assay was conducted with phages that formed CL and SCL to confirm their host spectrum.

### Random Amplification of Polymorphic DNA (RAPD)-PCR Subtyping of Phages

RAPD-PCR is used to genetically differentiate phage isolates. The analysis was performed using AccuPower PCR PreMix, and a mixture of primers OPL5 (5’-ACGCAGGCAC-3’), P1 (5’-CCGCAGCCAA-3’), and P2 (5’-AACGGGGCAGA-3’) (Lee, 2023). The following conditions were applied: initial denaturation at 94[for 5 min; 16 cycles of 94[for 45 s, 30[for 90 s, and 72[for 60 s; a second pre-denaturation at 94[for 3 min; 16 cycles of 94[for 30 s, 36[for 30, and 72[for 60 s; and a final extension at 72[for 10 min. The amplicon products were analyzed by electrophoresis. Ten microliters of the products were loaded onto a 1.5% agarose gel containing 0.02% EcoDye (SolGent Co, Ltd.). A 1 kb plus DNA ladder (SolGent Co., Ltd.) was used as the DNA marker. The products were electrophoresed in 1X Tris-Acetate-EDTA Buffer (iNtRON Biotechnology, Inc., Korea) at 100 V for 30 min. The gels were visualized using a Gel Doc system (ATTO, Korea).

### Whole Genome Sequencing (WGS) and Bioinformatics Analysis of 14 Selected Phages

Firstly, the genomic DNA of phages was extracted using the phenol-chloroform method (Texas A&M University, 2018). Five hundred microliters of phage solution were incubated for 1 h at 37[with 1.25 μL each of Dnase1 and RNase1. The solution was then incubated at 60[for 1 h after adding 20 μg of Proteinase K, 0.5% final concentration of 10% SDS stock, and 20 mM of 0.5 M EDTA (pH 8.0). An equal volume of phenol-chloroform was added, and the mixture was centrifuged at 6,000 rpm for 5 min at 25°C. The supernatant (approximately 450 μL) was carefully transferred to a fresh e-tube, and the phenol-chloroform extraction process was repeated once more. One-tenth of the total volume of 3M Sodium Acetate (pH 7.5) and 2.5 times the volume of 100% ethanol was added. The mixture was incubated overnight at -20°C. The DNA was resuspended in 50 μL of RNase-free water and then measured by using a NanoDrop spectrophotometer. The genomic DNA of phages was sequenced using an Illumina Miseq platform (San Diego, CA, USA), and the sequencing reads were assembled using the Celemics BTSeq analysis pipeline (v.2.27. Celemics, Korea).

The assembled sequences and open reading frames (ORFs) were predicted using Glimmer 3.02 and the Geneious Prime^®^ (v.2023.0.1. Biomatters Ltd.) software package. The predicted ORFs were annotated using BLASTP against the NCBI non-redundant protein database. Additionally, Geneious Prime was used to visualize and manually curate the annotations.

Toxin gene analysis was conducted by annotating the genomic data using the RAST server (Retrieved from https://rast.nmpdr.org/rast.cgi?page=Jobs), which identifies toxin genes based on its integrated database. Antibiotic resistance gene analysis was performed using ResFinder (Retrieved from https://cge.food.dtu.dk/services/ResFinder).

Phylogenetic analysis was conducted using the Virus Classification and Tree Building Online Resource (VICTOR. Retrieved from https://www.dsmz.de/) with the Genome-BLAST Distance Phylogeny method (GBDP). The phylogenetic tree was generated based on intergenomic distances calculated by the VICTOR tool. For proteomic analysis, the Viral Proteomic Tree (VipTree server. Retrieved from https://www.genome.jp/viptree/) web server was used, which calculates genome-wide sequence similarities using tBLASTx. The proteomic tree was <u>visualized</u> and annotated to show the relationships between the 14 phages and other known Klebsiella phages.

### Accession Numbers of The Phage Genome Data

The complete genome sequences for the isolated phages have been deposited in the NCBI database with the accession numbers: vB_KpnM_W1W2_10 (PP209575), vB_KpnM_W2-7 (PP273506), vB_KpnM_W7-3 (PP273507), vB_KpnM_W8-3 (PP273507), vB_KpnM_W8-4 (PP273509), vB_KpnM_W9-2 (PP312924), vB_KpnM_W10-2 (PP728205), vB_KpnM_W11-4 (PP312925), vB_KpnM_W12-3 (PP728206), vB_KpnM_W14-1 (PP312926), vB_KpnM_W14-2 (PP728207), vB_KpnM_W14-4 (PP312927), vB_KpnM_W15-2 (PP312928), vB_KpnP_W17 (PP728208).

### Morphological Observation of 14 Selected Phages

Ten microliters of phage lysate were applied to a copper grid for 2 min, and the excess liquid was removed with filter paper. The grids were then stained with 10 μL of 2% uranyl acetate for 30 s, and the excess liquid was absorbed. The grids were air-dried for 15 min before examining the phage morphology under a transmission electron microscope (TEM) at 35,000x magnification (Hitachi HT 7700 bio-TEM).

## Results

### The Characteristics of CRKP Clinical Isolates

A total of 24 isolates of *K. pneumoniae* were used in this study, comprising 23 clinical isolates and one ATCC strain. Table 1 details the phenotypic and genotypic characteristics of the CRKP clinical isolates. MLST analysis revealed that the 23 clinical isolates were distributed across 10 STs. The majority were identified as ST307 (n = 12) and ST11 (n = 4), both of which have recently emerged as high-risk clones of concern in Korea (Cho et al., 2022; Yoon et al., 2018). The remaining isolates were classified into individual STs: ST15, ST48, ST133, ST336, ST392, ST789, and novel ST. According to the allele profile, the novel sequence type KBN10P07305 has been registered in the PasteurMLST database as ST7145 (Table 1).

The results of the Carba NP test were interpreted based on color changes. A red-to-yellow or red-to-orange color change indicated the presence of carbapenemase production. All isolates were positive for carbapenemase activity except for three (KBN10P02349, KBN10P06655, and KBN10P07294).

The string test results revealed that three isolates (KBN10P05309, KBN10P06644, and KBN10P07146), along with the ATCC13883 strain, formed viscous strings longer than 5 mm, indicating positive results for HMV, which is associated with hypervirulent strains (Table 1).

A total of 10 distinct *wzi* alleles (3, 18, 19, 50, 62, 123, 141, 173, 178, and 180) were identified. The *wzi* allele number determined five capsular types: K3, K15 (including K17, K50, K51, and K52 variants), K18, K19, and K62. The remaining strains were classified according to their K-locus types, which included four types: KL25, KL102 (including KL149 and KL155 variants), KL116, and KL136. However, *wzi* allele 178 could not be associated with a specific K or K-locus type. All isolates were found to carry the KPC-2 gene, while a single isolate, KBN10P06378, was identified as carrying the OXA-48 gene, as shown in Fig. S1.

### Isolation of Phages

A total of 38 phages were isolated using four strains of *K. pneumoniae* (KCTC 2208, ATCC 13883, clinical isolate KBN10P05309, or KBN10P07398) as hosts. These phages were named according to the phage nomenclature rules (Adriaenssens and Brister, 2017). For example, in the name vB_KpnM_W2-7, “W” stands for sewage, the number “2” indicates the sewage sample number, and “7” represents the isolation sequence of the phage from that sample. Information regarding the phage isolates and their titers is presented in Table 3. Examination of plaque morphology revealed that vB-_KpnP_W17 formed a significant halo (Fig. 1).

**Fig. 1.**
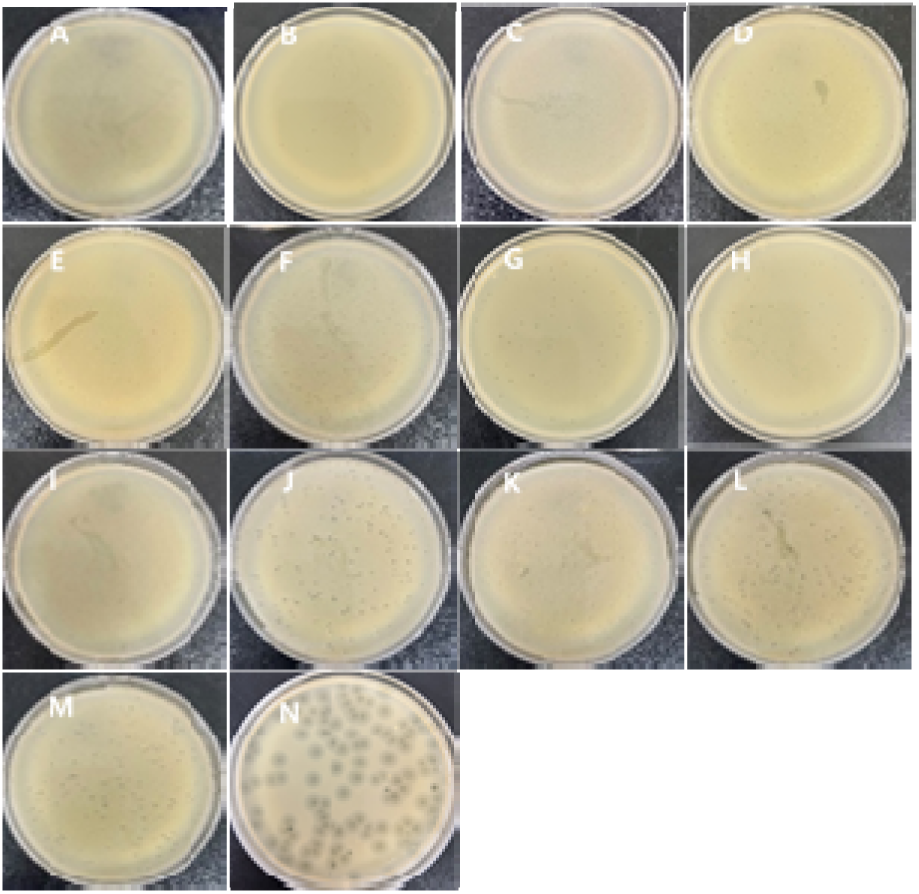
Plaque morphology of *Klebsiella pneumoniae* phages. (A) vB_KpnM_W1W2-10 (B) vB_KpnM_W2-7 (C) vB_KpnM_W7-3 (D) vB_KpnM_W8-3 (E) vB_KpnM_W8-4 (F) vB_KpnM_W9-2 (G) vB_KpnM_W10-2 (H) vB_KpnM_W11-4 (I) vB_KpnM_W12-3 (J) vB_KpnM_W14-1 (K) vB_KpnM_W14-2 (L) vB_KpnM_W14-4 (M) vB_KpnM_W15-2 (N) vB_KpnP_W17

### Lytic Spectrum of Phage Isolates

The lytic spectra of 38 phages were determined using 23 CRKP isolates and ATCC 13883 strain. Most phages lysed the ST11 CRKP isolates (Table 2). Although the original host for phages vB_KpnM_W1W2-1, vB_KpnM_W1W2-10, vB_KpnM_W2-7, vB_KpnM_W2-8, and vB_KpnM_W2-9 was KCTC 2208, these phages were cultured using the clinical isolate KBN10P05309 as a host due to ineffective propagation in the original host. No phages were isolated when clinical isolates KBN10P02346 and KBN10P02349 were used as hosts, and none of the isolated phages lysed KBN10P02346. The isolate KBN10P07139 was infected by most phages, with phage vB_KpnM_W10-4 exhibiting the widest lytic spectrum. Phages lysed strains belonging to the same STs as their respective hosts. Phages using KBN10P05309 as the host lysed all ST11 strains, while those using KBN10P07398 as the host lysed most ST307 strains. Since the spot assay does not provide detailed information about phage infection, we conducted plaque assays using phages that exhibited clear and turbid lysis zones in the lytic spectrum. The efficiency of plating (EOP) results indicated intermediate EOP values (>0.1-1) for strains of the same STs except for the host strain. However, strains from different STs displayed low EOP values and could not form plaques, failing to obtain EOP values (Table 2).

**Table 2.**
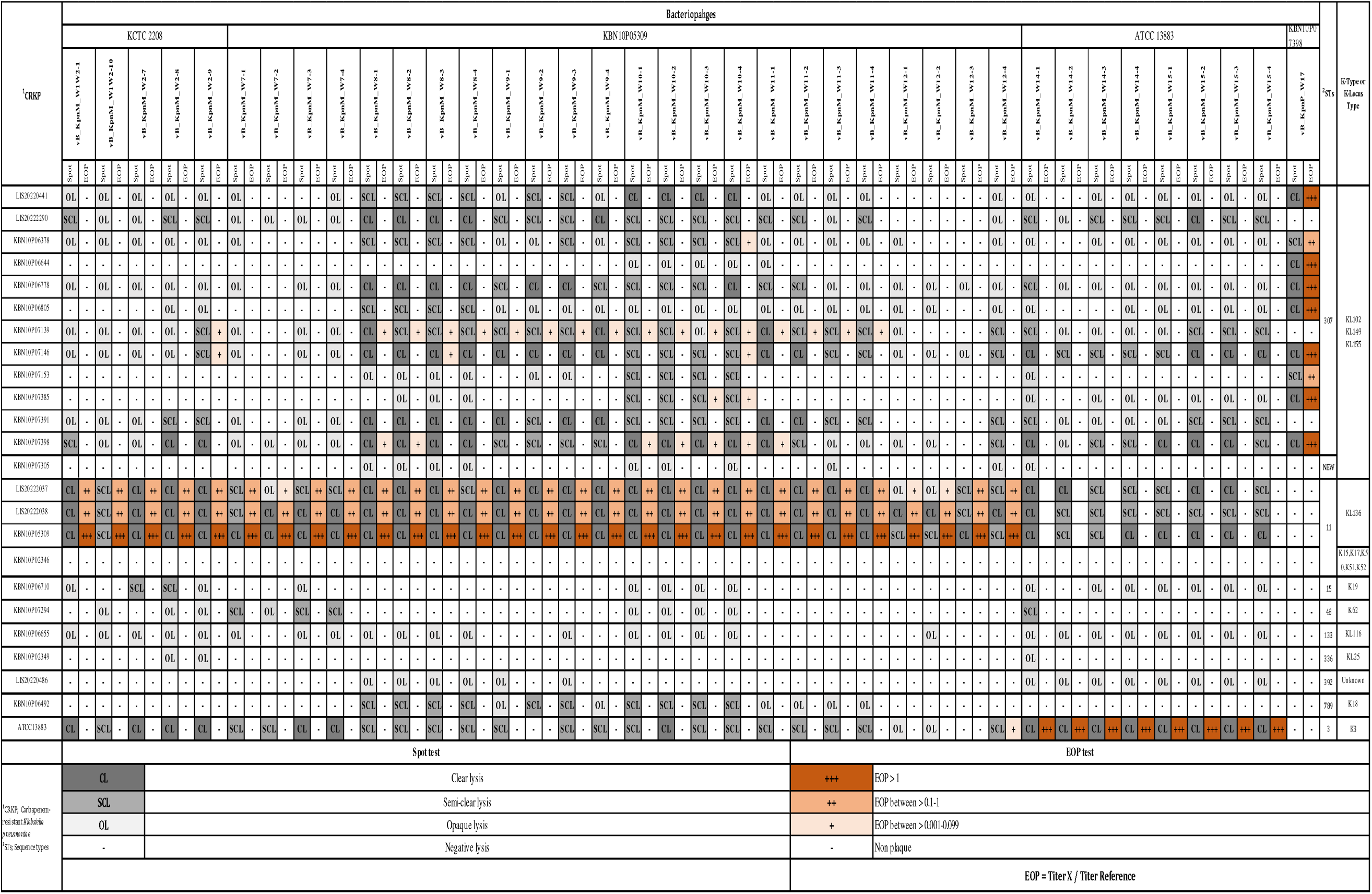
Phage lytic spectrum and efficiency of plating against various Klebsiella pneumonaie ST types.

### Subtypes of Phage Isolates

The sizes of the RAPD amplicons ranged from approximately 200 bp to 3 kb. Based on the band patterns, phages were initially classified into five groups. One to two phages were chosen from each group for comprehensive genomic analysis, considering both their band patterns and lytic spectra. Specifically, phages were first categorized according to their lytic spectra. Then, RAPD-PCR results were used to select 14 phages that showed distinct differences, ensuring representation of the diversity within the groups. These 14 phages were chosen for further WGS study (Fig. 2).

**Fig. 2.**
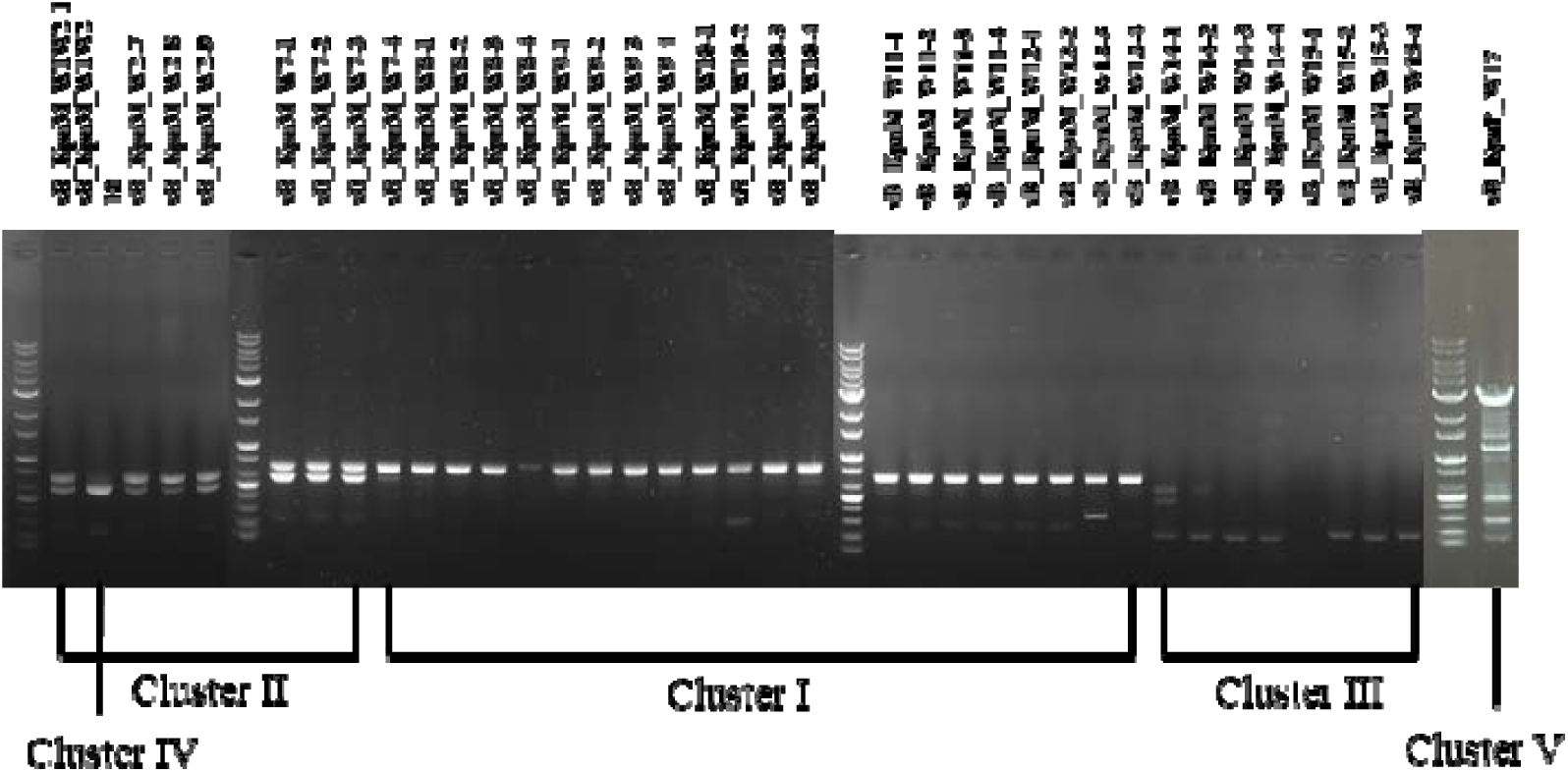
Representative RAPD gel image.

### Morphological Characterization of The Phages

TEM observations showed that 13 phages possessed long contractile tails, base plates with short spikes, and tail fibers, identifying them as *Myovirus* morphotypes. The other one phage exhibited the distinct structure of a *Podovirus* morphotype, featuring a small, round icosahedral head and a short tail. Representative TEM images of the 14 phages are shown in Fig. 3. The length information for each phage is specified in Table 3.

**Fig. 3.**
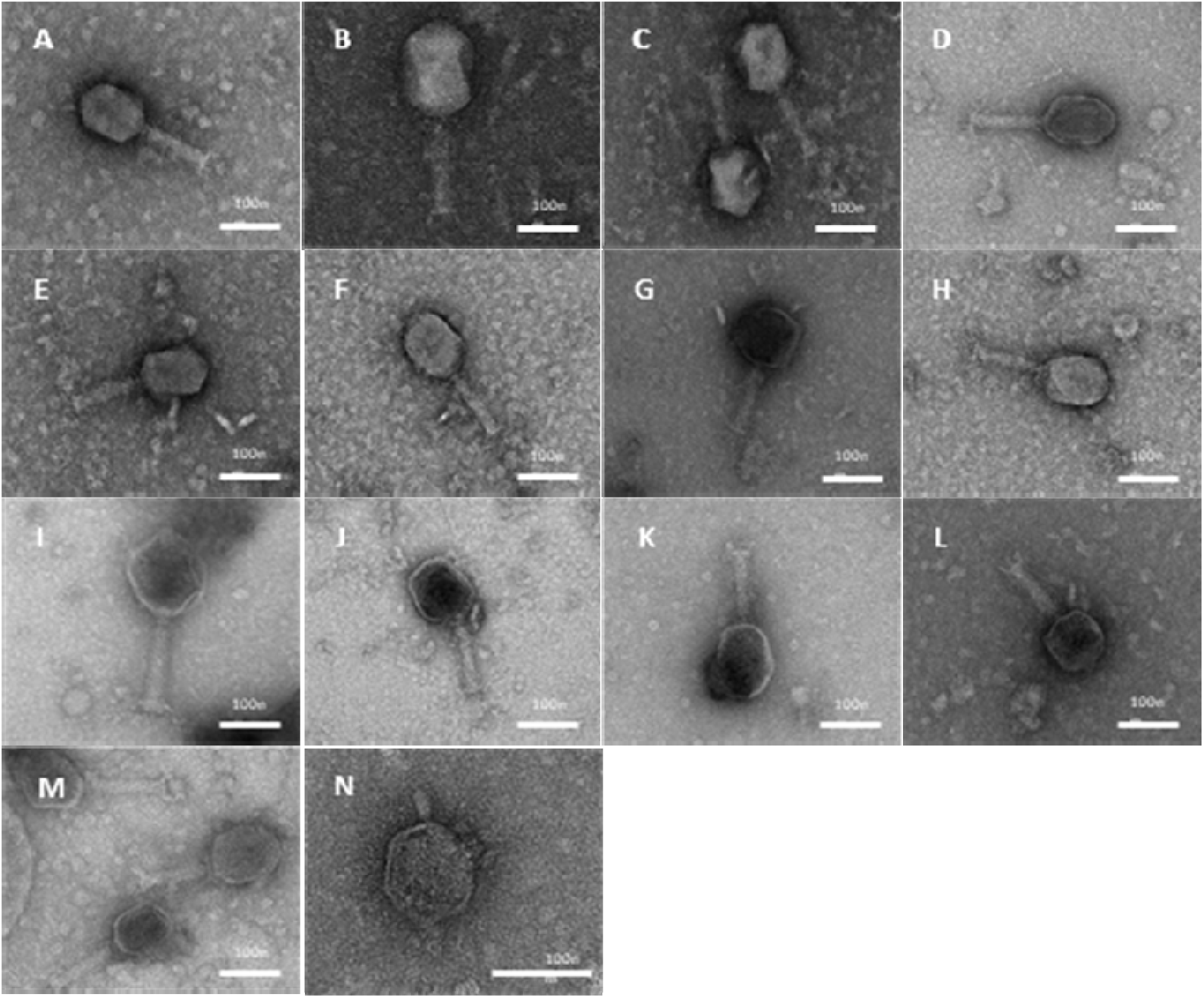
Transmission electron microscopy images of *Klebsiella pneumoniae* lytic phages. (A) vB_KpnM_W1W2-10 (B) vB_KpnM_W2-7 (C) vB_KpnM_W7-3 (D) vB_KpnM_W8-3 (E) vB_KpnM_W8-4 (F) vB_KpnM_W9-2 (G) vB_KpnM_W10-2 (H) vB_KpnM_W11-4 (I) vB_KpnM_W12-3 (J) vB_KpnM_W14-1 (K) vB_KpnM_W14-2 (L) vB_KpnM_W14-4 (M) vB_KpnM_W15-2 (N) vB_KpnP_W17

**Table 3.**
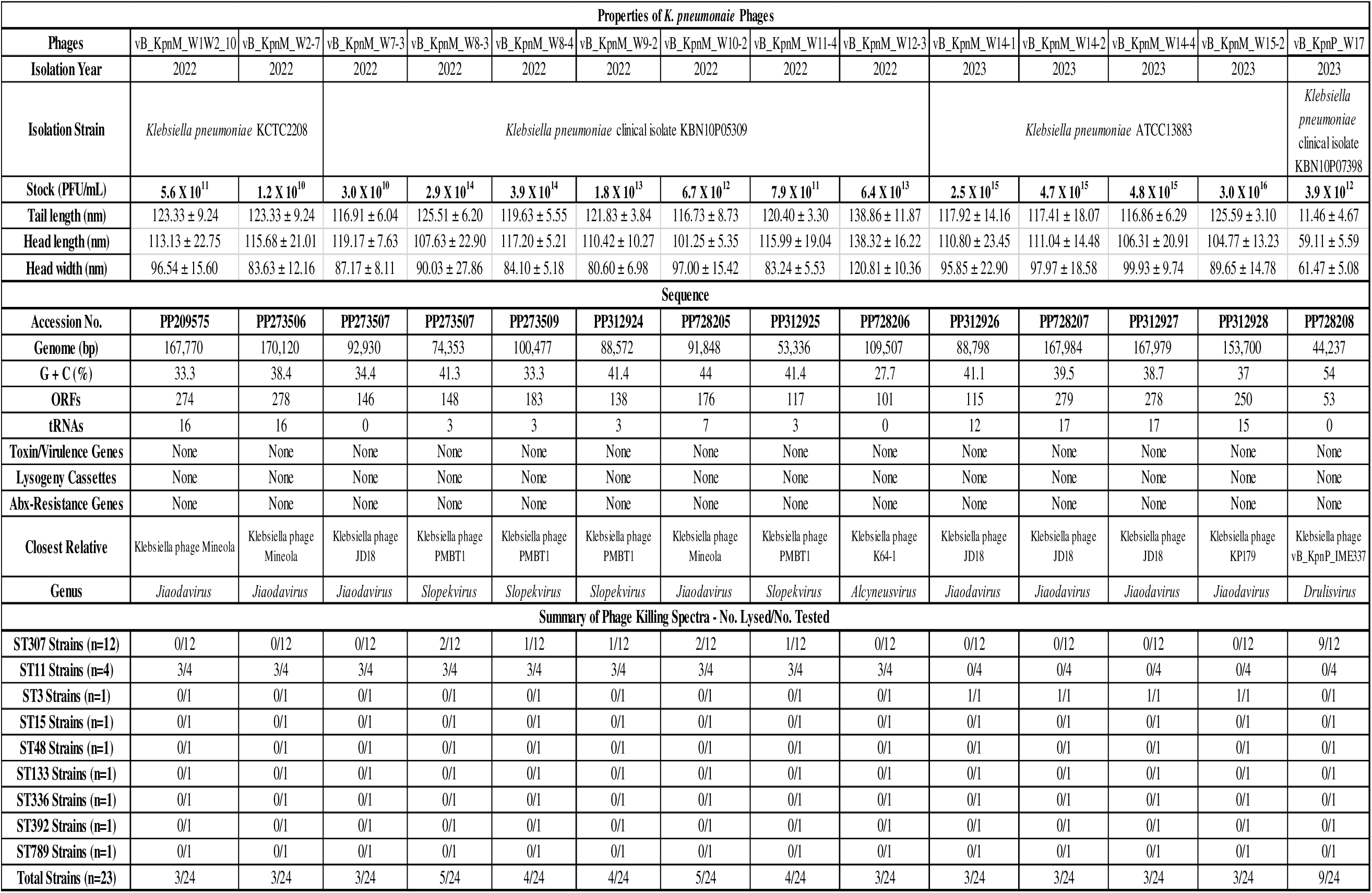
Summary of characterization of Klebsiella pneumoniae phages.

### Genomic Characterization of the Phages

A total of 14 phages were genomically characterized. Of these, eight phages were identified as belonging to the *Jiaodavirus* genus, and four phages were classified under the *Slopekvirus* genus, both of which are part of the *Straboviridae* family. The phage vB_KpnM_W12-3 was found to be a member of the *Alcyneusvirus* genus, although its family classification remains unknown. The phage vB_KpnP_W17 was classified under the *Drulisvirus* genus, part of the *Autographiviridae* family. The phages had genome sizes ranging from 44 kb ∼ 170 kb and G+C contents between 27.7% ∼ 54.0%. The isolated phages showed high similarity to *Klebsiella* phage Mineola (vB_KpnM_W1W2-10, vB_KpnM_W2-7), *Klebsiella* phage JD18 (vB_KpnM_W7-3, vB_KpnM_W14-1, vB_KpnM_W14-2, vB_KpnM_W14-4), *Klebsiella* phage PMBT1 (vB_KpnM_W8-3, vB_KpnM_W8-4, vB_KpnM_W9-2, vB_KpnM_W11-4), *Klebsiella* phage K64-1 (vB_KpnM_W12-3), *Klebsiella* phage KP179 (vB_KpnM_W15-2), and *Klebsiella* phage vB_KpnP_IME337 (vB_KpnP_W17) with identities ranging from 88% to 99.15%. The genomes of these phages encode between 53 and 279 ORFs, with approximately 50% being hypothetical proteins and the remaining 50% functional genes. The coding sequences (CDS) annotation was conducted with over 90% similarity. Functional genes were divided into four groups based on their roles in the phage infection cycle: translation and transcription-related proteins, structural proteins, endolysin-related proteins, and others (Fig. 5). No bacterial toxin genes, antibiotic resistance genes, or temperate life cycle genes were found in the genomes (Table 3, Fig. 4). A proteomic tree was constructed to include all *Klebsiella* phages and those isolated in this study, while a phylogenetic tree was explicitly built for the 14 phages isolated in this study. The proteomic and phylogenetic trees indicate that the 14 phages were classified into four groups (Fig. 4).

**Fig. 4.**
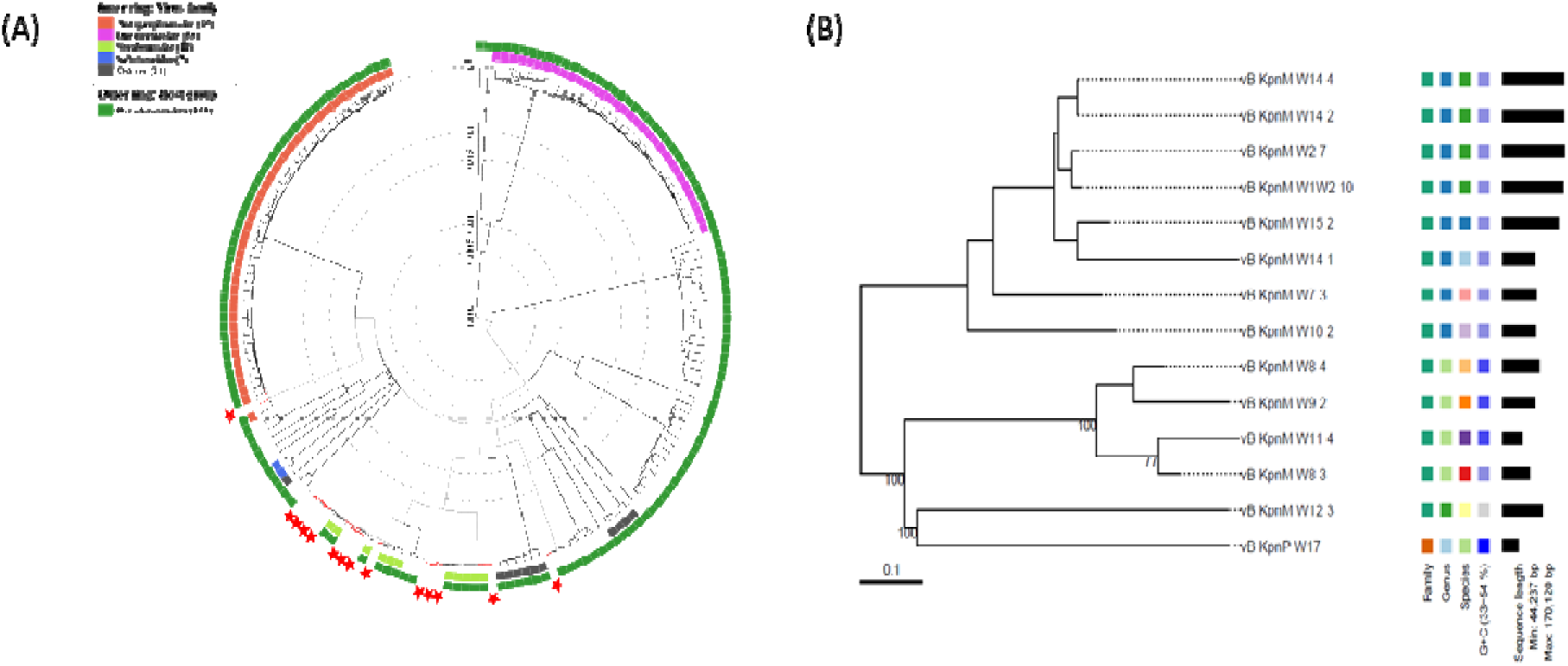
Dendrogram of selected phages based on genomic DNA sequences. (A) is the proteomic tree of Klebsiella pneumoniae phages. Red stars indicate 14 isolated phages. (B) is the phylogenetic tree of *Klebsiella pneumoniae* phages.

**Fig 5.**
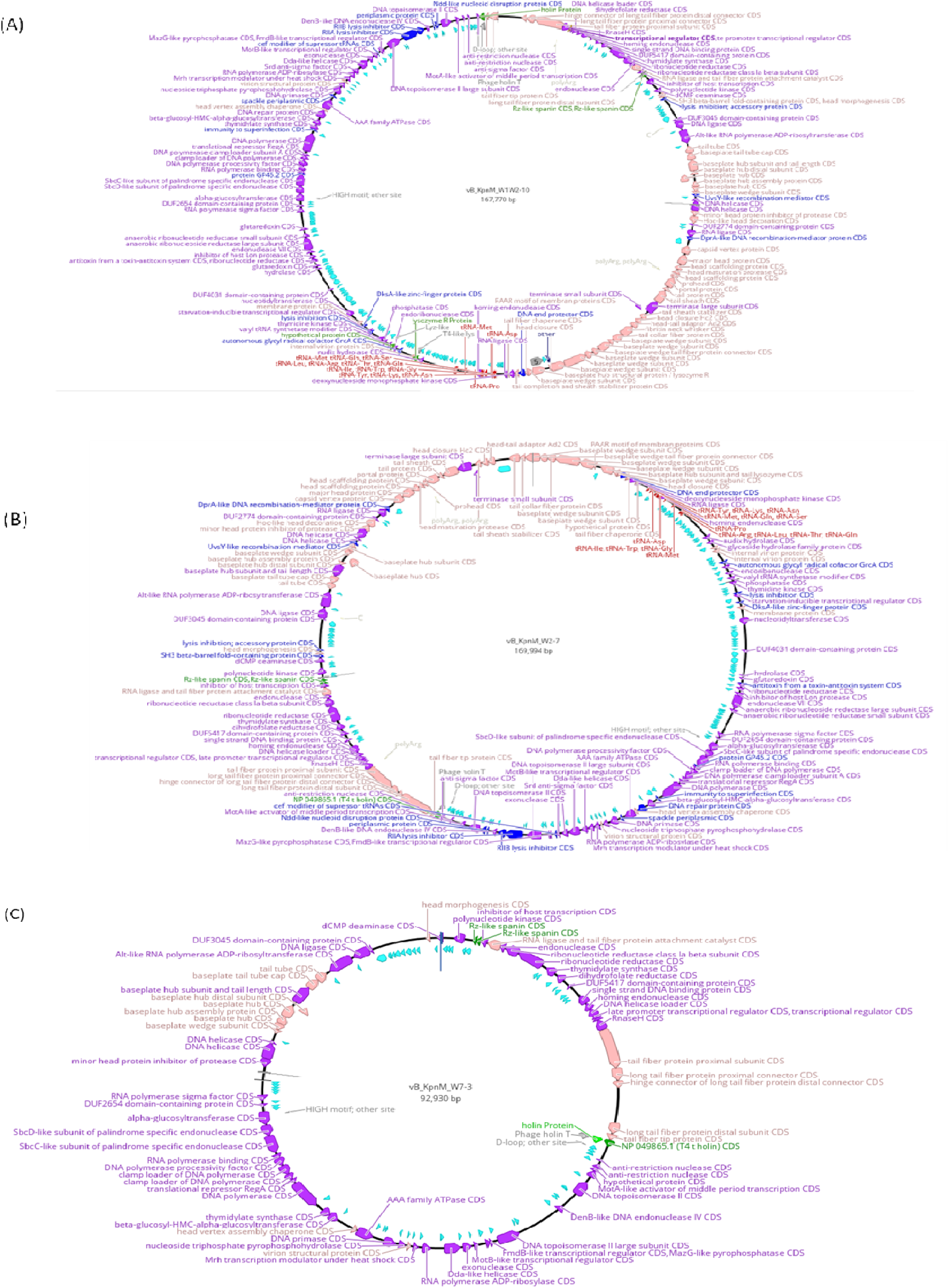
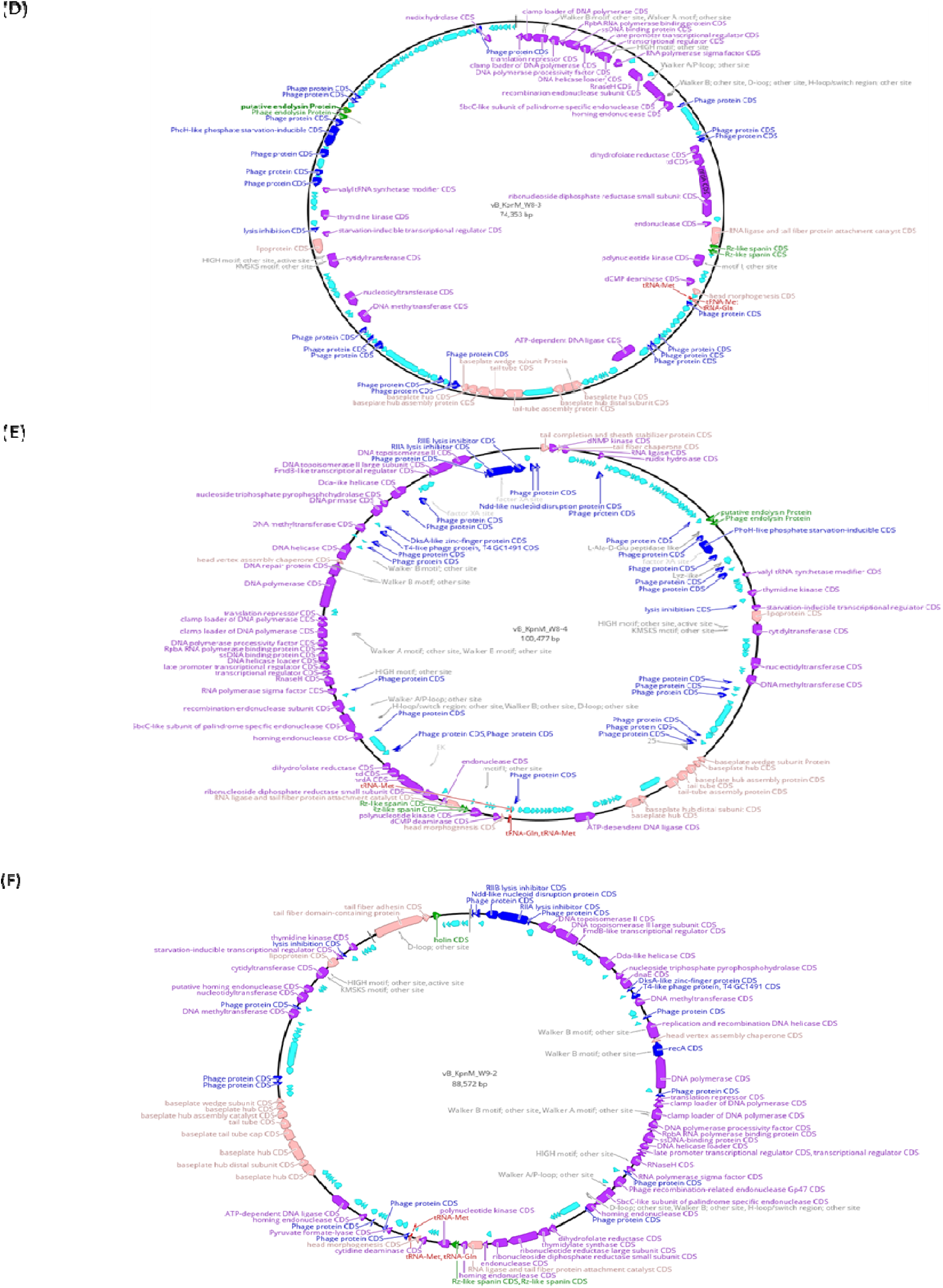
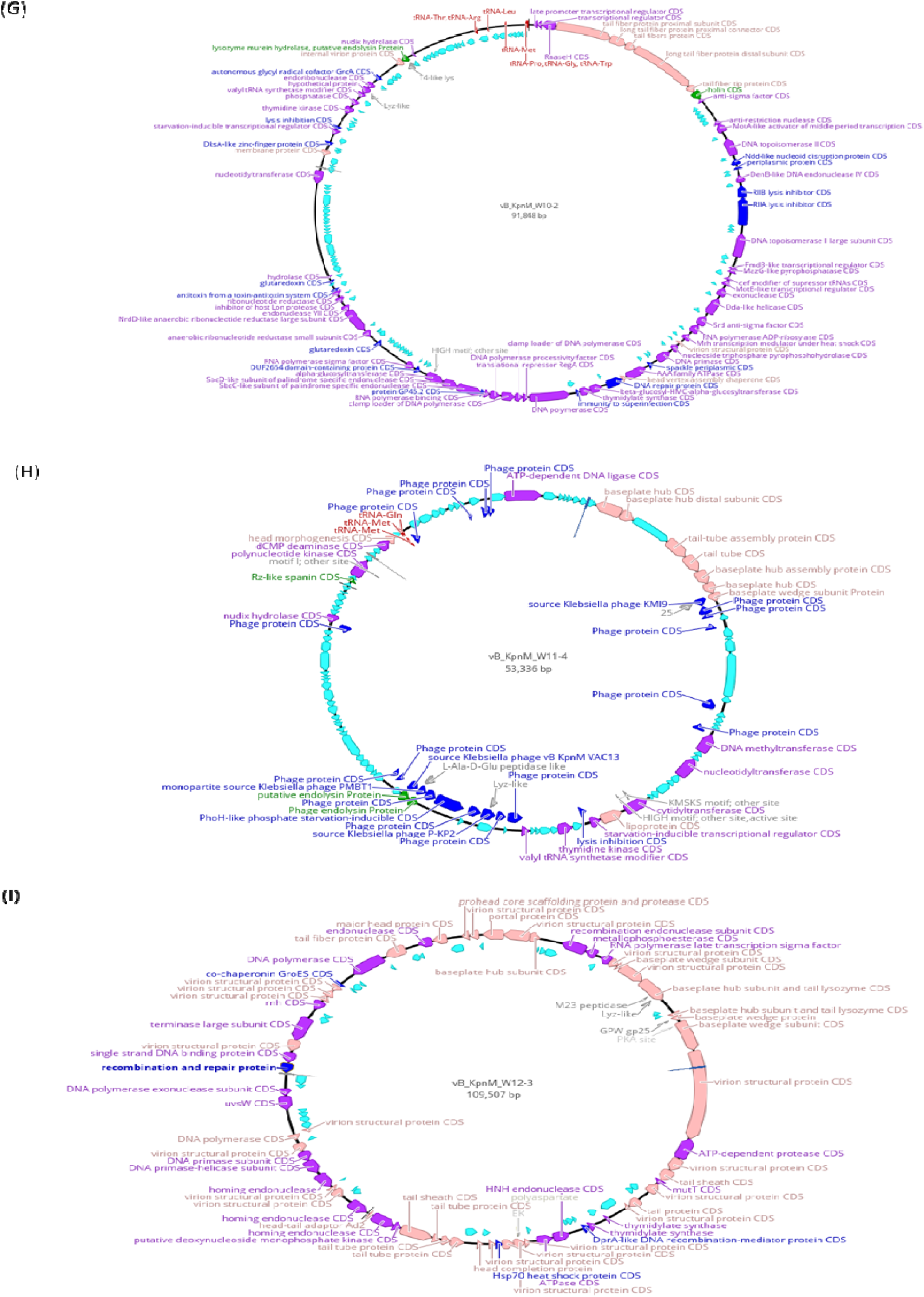
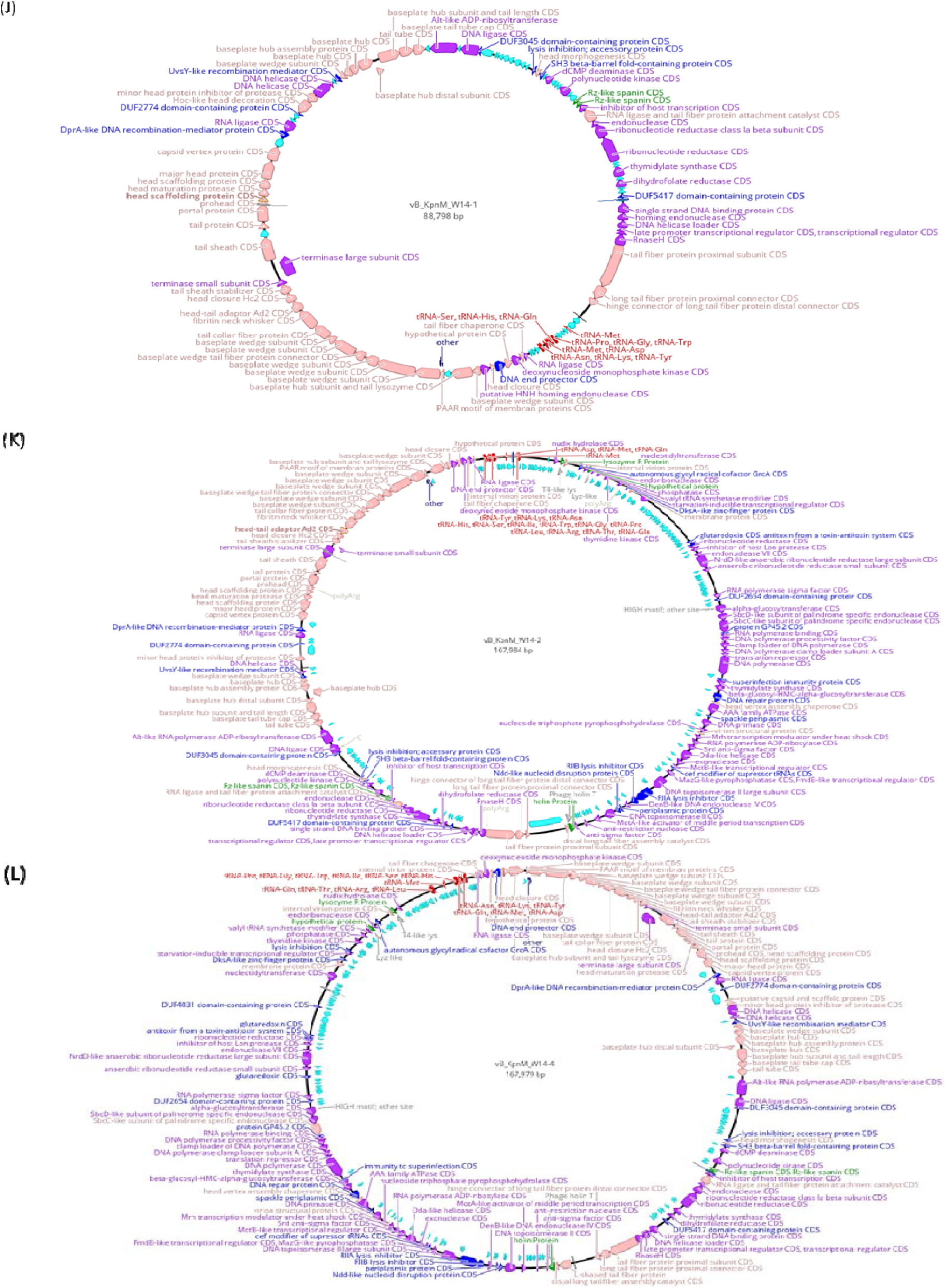
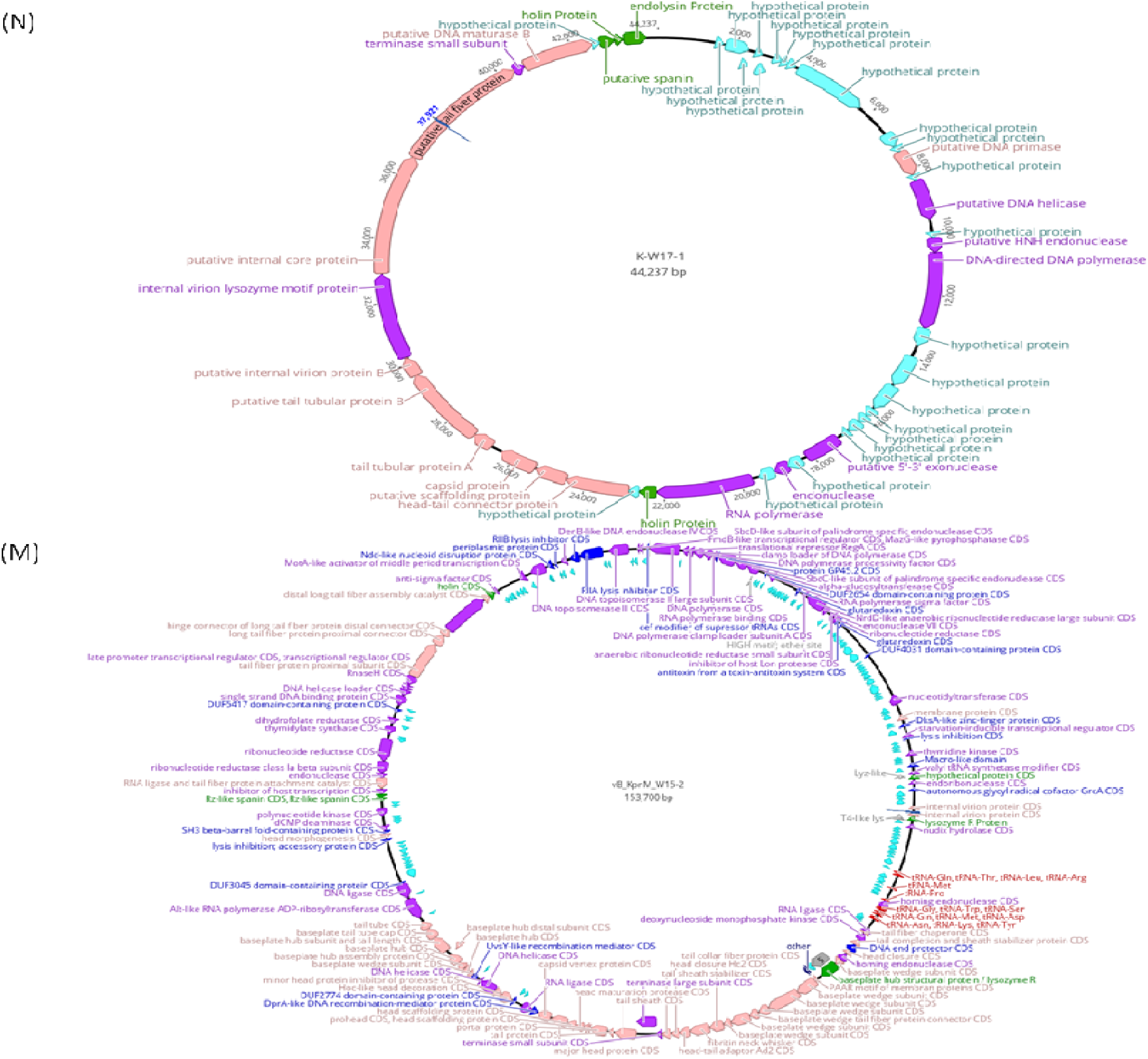
Genome maps of 14 selected phages. CDS and transcription direction are indicated by arrows. Each color of the arrow represents the function of CDS. Purple: Translation and transcription-related protein; Pink: Structural protein; Green: Endolysin-related protein; Skyblue: Hypothetical protein; Blue: Others (Protein that does not belong to the above four categories)

## Discussion

This study aimed to develop effective phage therapy against CRKP, a major member of the ESKAPE group of pathogens (*Enterococcus faecium, Staphylococcus aureus, Klebsiella pneumoniae, Acinetobacter baumannii, Pseudomonas aeruginosa, and Enterobacter species*) known for their antibiotic resistance (Moellering, 2010; Zurabov and Zhilenkov, 2021). To ensure the success of phage therapy against *K. pneumoniae* infections, it is crucial to thoroughly understand the diverse phenotypic and genotypic characteristics of both *K. pneumoniae* pathogens and their associated phages (Li et al., 2021). In this study, we successfully characterized 23 CRKP clinical isolates. We analyzed 38 phage isolates obtained from sewage samples, revealing several key insights that could inform the development of effective phage therapy against CRKP.

Initially, KCTC 2208 was used as the host for phage isolation. Although phage isolation was initially successful, amplification did not occur when attempting to increase the titer. Consequently, all phages originally isolated using KCTC 2208 were transitioned to KBN10P05309 as the host for further studies.

None of the phages lysed KBN10P02346, and no specific phages could be isolated for KBN10P02346 and KBN10P02349. While the other CRKP clinical isolates were collected after 2017, these two were isolated in 2011, which may explain this phenomenon, although further analysis is required.

Although phages vB_KpnM_W14-2 and vB_KpnM_W14-4 were initially considered different based on phage classification, sequencing results reveal a 99.72% similarity, with only a minor difference of five nucleotides, suggesting that they are likely the same phage.

According to the results of *wzi* gene sequencing, the isolates had 10 different K-type and K-locus types. Since capsular polysaccharides (CPS) are the most important virulence factors of *K. pneumoniae* and interfere with phage attachment to the cell, understanding these characteristics is essential for phage therapy (Horváth et al., 2020; Zurabov and Zhilenkov, 2021).

The presence of depolymerase enzymes in phages is particularly noteworthy, as these enzymes can degrade the CPS of *K. pneumoniae*, facilitating phage attachment and subsequent bacterial lysis (Pan et al., 2017). The appearance of a halo—a turbid zone surrounding a plaque—indicates depolymerase activity, which can rapidly identify capsule-targeting phages (Knecht et al., 2020). In the case of phage vB_KpnP_W17, which belongs to the *Podoviridae* family, a significant double halo was observed**-** a characteristic commonly associated with *podoviruses*. This phage exhibited halo formation morphologically and showed the presence of depolymerase genes, specifically in components like the tail fiber and baseplate. Moreover, genomic analysis of several phages revealed the presence of three types of lysozymes, including baseplate hub subunits and tail lysozymes, all predicted to have depolymerase activity. Since *K. pneumoniae* is known for its high capsule production, discovering these lysozymes with potential capsule-removing capabilities marks a significant step towards overcoming bacterial defenses. These characteristics are illustrated in Fig. 1 and Fig. 5.

Our findings highlight the ST-specific lytic activity of several isolated phages. In host spectrum determination, spot assay results showed that the phages broadly lysed the isolates from various STs. However, in the plaque assay, phage plaques were predominantly observed in clinical isolates belonging to the same STs as the host, with only a few plaques formed in isolates of different STs due to limited infections. This suggests that while phage depolymerase activity was effective in causing bacterial lysis, it fell short of achieving complete infection. This ST-specific activity implies a strong correlation between the phage host range and the bacterial strain’s genetic background, supporting the potential for developing tailored phage cocktails. However, this specificity also presents a limitation: the narrow host range may reduce the applicability of individual phages across diverse bacterial populations. Therefore, a broad and diverse phage library will ensure comprehensive coverage and effective treatment of CRKP infections in varied clinical settings. Moreover, the effectiveness of phages was confined to *K. pneumoniae*, with no lytic activity observed against other ESKAPE pathogens in the spot test, indicating species-specific actions.

Beyond these findings, our study contributes to the ongoing efforts to categorize phages based on their susceptibility to clinical isolates according to STs. This approach aims to adapt isolated phages for therapeutic use by better understanding the mechanisms of phage infection. The consistency of our results with this categorization method supports the potential for tailoring phage therapy to specific bacterial strains.

We plan to expand our research by acquiring WGS data for a more significant number of bacterial strains and phages. This expanded dataset will enable us to develop a program capable of predicting phage infection mechanisms, further enhancing the precision and effectiveness of phage therapy. By integrating genomic data into our analysis, we aim to improve our understanding of how phages interact with their bacterial hosts and ultimately optimize phage selection for therapeutic applications.

Despite the promising advancements in our study, several challenges must be addressed before phage therapy can be widely implemented. A significant gap remains in the availability of well-characterized phages against *K. pneumoniae*, highlighting the need for continued isolation and characterization of a broader range of phages. This is particularly important given the narrow host range observed in some phages, which could not lyse certain CRKP strains, underscoring the necessity of expanding the phage library to improve the versatility of phage therapy. Additionally, although our study successfully identified phages with lytic activity, the efficacy of these phages *in vivo* remains to be validated. *In vivo* studies are crucial to assess the therapeutic potential of these phages in real-world clinical settings, where factors such as the host immune response and the complex microbial environment may influence treatment outcomes (Górski et al., 2023).

To mitigate bacterial resistance, one effective strategy is using phage cocktails, which combine multiple phages targeting different bacterial receptors (Mateus et al., 2014; Hasan and Ahn, 2022). Another promising strategy is using phage-antibiotic combinations, which offer a synergistic effect by enhancing antibiotic effectiveness and increasing bacterial susceptibility to phages (Gu et al., 2020; Hasan and Ahn, 2022; Choi et al., 2024b). This dual approach not only boosts bactericidal effects but also makes it more difficult for bacteria to develop resistance to both treatments simultaneously (Mateus et al., 2014; Gu et al., 2020; Hasan and Ahn, 2022). Furthermore, there is growing interest in the adaptive design of phages, where phages are engineered to evolve alongside bacterial mutations, potentially extending the utility of phage therapy by ensuring its continued effectiveness even as bacterial populations change (Gencay et al., 2024; Salmond and Fineran 2015).

In conclusion, this study provides valuable insights into the development of phage therapy as a viable alternative to traditional antibiotics for treating infections caused by CRKP. The specificity of phage action against certain STs, coupled with the challenges of broadening the host range, defines the current limits of our understanding and the scope for future investigation. As such, ongoing research and clinical trials are essential to fully harness the potential of phages in addressing the challenges posed by antibiotic resistance.

## Acknowledgment(s)

This research was funded by a grant from the Korea Disease Control and Prevention Agency (Grant No. 2022-ER2202-00).

## Figures

**Fig. S1.**
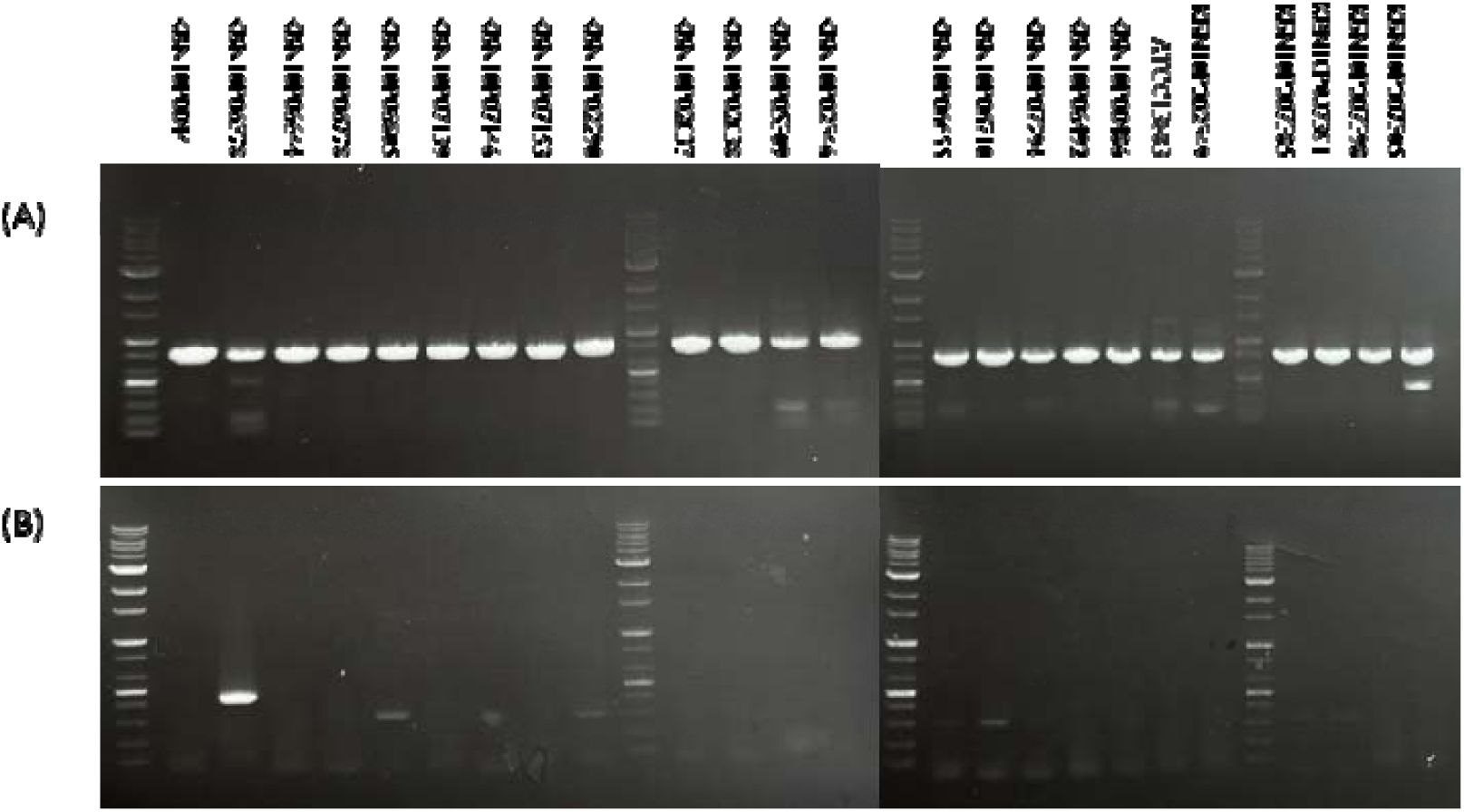
Agarose gel images of multiplex PCR for carbapenemase genes detection in carbapenem-resistant *Klebsiella pneumoniae*. (A) *bla*_KPC-2_ (amplicon size 798 bp), (B) *bla*_OXA-48_ (amplicon size 438bp).

